# From Family Relationship to Regulation: Parent–Child Amygdala Activation Similarity is Linked to Prefrontal Recruitment and Youth Emotional Adaptation

**DOI:** 10.1101/2025.07.21.665810

**Authors:** Ya-Yun Chen, Zexi Zhou, Yang Qu, Tae-Ho Lee

**Affiliations:** Department of Psychology, Virginia Tech, Blacksburg, VA, USA; Department of Human Development and Family Sciences, The University of Texas at Austin, Austin, TX, USA; School of Education and Social Policy, Northwestern University, Evanston, IL, USA; School of Neuroscience, Virginia Tech, Blacksburg, VA, USA

**Keywords:** parent–child neural similarity, family relationship, emotional adaptation, amygdala-prefrontal connectivity

## Abstract

Extensive research highlights the importance of parent–child similarity at psychological, behavioral, and physiological levels in fostering children’s adaptation. While emerging studies have begun to explore neural similarity, little is known about the factors that influence parent– child neural similarity during adolescence or how such similarity may support youth emotional adaptation when coping with heightened uncertainty. The current study investigated the relations among family relationship quality, parent–youth neural similarity, and youth emotional adaptation using task-based fMRI. With 50 participants (25 parent–youth dyads; parent: *M*_age_ = 43.88±7.42 years, 72% female; youths: *M*_age_ = 11.16±2.85 years, 44% female), we found that higher parent–youth neural similarity in the amygdala activation patterns during emotionally uncertain situations was associated with better youths’ emotional adaptation, as indicated by lower levels of anxiety and depression and higher resilience. This neural similarity mediated the beneficial effects of positive family relationships on youths’ emotional adaptation. Additionally, amygdala activation similarity was associated with amygdala to dorsolateral prefrontal cortex (dlPFC) connectivity during uncertainty in youth, revealed by generalized psychophysiological interaction (gPPI) analyses. This amygdala-dlPFC connectivity, in turn, was linked to better emotional adaptation in youth. Importantly, these findings were specific to uncertainty-related bottom-up processing, rather than effortful top-down regulation processing or responses to certain aversive stimuli. These findings offer new insight into how relational and neural factors jointly shape youth emotional adaptation, and suggest that parent–child neural similarity in the amygdala may serve as a mechanism for supporting implicit emotion regulation, helping youth cope with uncertain situations.

**Significance Statement:** During adolescence, youth begin to expand their social worlds beyond the family, yet family relationships continue to shape how they navigate emotionally complex environments. How do these relationships influence youth brain function and emotional well-being? Using task-based fMRI, we found that greater parent–child neural similarity in the amygdala during emotionally uncertain situations is associated with youths’ lower anxiety and depression and greater resilience. This amygdala neural similarity was also linked to prefrontal engagement in youth during early arousal stage, supporting adaptive implicit regulation under uncertainty. These findings suggest that neural alignment between parents and youths may serve as a key mechanism by which positive family relationships promote emotional adaptation, highlighting the enduring role of family context in adolescent brain development.

## Introduction

Identifying a potential neural mechanism that supports adolescents’ emotional adaptation is crucial during this critical transitional period, which requires them to navigate complex social and emotional challenges, including stress related to uncertainty about their environment and future (Murberg and Bru, 2004; Byrne et al., 2007; Lim and Javadpour, 2021). Prior research suggests that greater neural similarity between parents and children may promote more adaptive outcomes during this period (Bell, 2020; Rogers et al., 2022; Qu et al., 2023). Therefore, the current study investigates whether and how parent–child neural similarity during uncertainty processing is associated with youth emotional adaptation.

Heightened sensitivity to stress and uncertainty in youth is associated with altered activity in their brain’s affective limbic circuit (Casey et al., 2008). The amygdala, a central structure within this circuit, is activated by a range of aversive stimuli (e.g., Nitschke et al., 2006; Janak and Tye, 2015; see review by Davis and Whalen, 2001), as well as in contexts involving uncertainty (e.g., Rosen and Donley, 2006; Belova et al., 2007; Herry et al., 2007; Sarinopoulos et al., 2010; Grupe and Nitschke, 2013). Amygdala alterations have been concurrently and prospectively linked to youth emotional dysregulation and increased risk for anxiety and depression (Williams et al., 2015; Albaugh et al., 2017; Tang et al., 2018; Strawn et al., 2021). Therefore, the amygdala may serve as a key neural region to examine the association between parent–child neural activation similarity during uncertainty processing and youth emotional adaptation.

In addition to examining amygdala as a single region, dual-systems model on adolescent brain development suggests that heightened emotional reactivity and difficulties in regulation during adolescence stem from asynchronous development between subcortical affective systems and prefrontal regulatory systems (Casey et al., 2008; Steinberg, 2008). In other words, successful emotional development and coping depend on balanced interactions between bottom-up affective processes and top-down regulatory mechanisms (McClure et al., 2004;

Steinberg, 2008; Casey, 2015). If the top-down system effectively regulates bottom-up emotional reactivity, this reflects a more mature and adaptive neurodevelopmental profile. Empirical studies have shown that amygdala-frontal connectivity is linked to stress responses and coping in youth (Gee and Casey, 2015). A meta-analysis further demonstrated that amygdala-frontal connectivity is robustly associated with successful emotion regulation (Berboth and Morawetz, 2021). Building on this evidence, if parent–child amygdala activation similarity during uncertainty processing is associated with youth emotional adaptation, the current study further examines whether this neural similarity is related to youth’s prefrontal engagement during uncertain situations, aiming to address a mechanistic question about how shared affective processing between parent and child may support regulatory function in youth.

Moreover, it is important to consider how such parent–child neural similarity emerges within the broader family context. Developmental neuroscience theories suggest that parent-child relationship quality may shape this similarity and, in turn, youth’s emotional development (Birk et al., 2022; Qu et al., 2023). High-quality family relationship, characterized by high cohesion (i.e., closeness and support), low conflict (i.e., strains and disagreement), and greater family identity (i.e., a sense of self shaped by connection to family), can foster shared dyadic processes that promote coordination and attunement (Fosco and Lydon-Staley, 2020; Cahill et al., 2021; Zhou et al., 2023), supporting youth’s emotional well-being (Ratliff et al., 2022). While prior research links positive family relationships to greater neural similarity under stress (Lee et al., 2018; Qu et al., 2023), no studies have examined the comprehensive process from family relationship to parent-youth neural similarity, and in turn, youth’s emotional adaptation.

Taken together, we hypothesized that greater parent–child amygdala activation similarity during uncertainty would be linked to better youth emotional adaptation, with positive family relationships predicting greater neural similarity. We further expected this neural similarity to mediate the effect of family relationship on emotional adaptation in youth, and to be associated with increased amygdala-prefrontal connectivity during uncertainty in youth, reflecting a regulatory mechanism that underlies the protective effect of parent–child neural similarity.

## Materials and Methods

### Participants and Procedure

Participants were recruited through Facebook group flyers, newspaper advertisements, and local media in the New River Valley area, Virginia, without restrictions on gender or race/ethnicity. Written informed consent was obtained, with children providing additional parental consent. The study protocol was approved by the Institutional Review Board of Virginia Tech. Exclusion criteria included MRI safety concerns, claustrophobia, significant head injury, orthodontia issues, severe psychopathology, and other MRI contraindications, assessed via self-report with parental assistance for children.

Initially, 32 parent-youth dyads were recruited (Total *N* = 64). However, one parent and two children discontinued the scan while in the scanner, two children had dental conditions affecting image quality, one child had unusual brain structures, and one parent had incomplete data, resulting in seven dyads excluded. The final sample included 25 parent-youth dyads (parents: *M*_age_ = 43.88 ± 7.42 years, 72% female; youth: *M*_age_ = 11.16 ± 2.85 years, 44% female). Parents self-identified as primary caregivers, with 88% being biological, 8% adoptive, and 4% guardians. Regarding race and ethnicity, 72% of youth identified as non-Hispanic White American, 12% as more than one race, 8% as non-Hispanic Asian American, 4% as Hispanic American, and 4% as non-Hispanic Black or African American. Among parents, 84% identified as non-Hispanic White American, 8% as non-Hispanic Asian American, 4% as non-Hispanic Black or African American, and 4% as more than one race.

Youth completed self-report measures of emotional adaptation (i.e., depression, anxiety, and ego resilience) and parent–child relationship quality prior to the MRI scan. Both youth and their parents then underwent three sessions of the Emotion Regulation Task during fMRI, along with T1- and T2-weighted structural imaging.

## Experimental Design

### Emotion Regulation Task

Participants completed the modified Emotion Regulation Task during fMRI scanning. Each trial consisted of four phases: cue presentation (2-3 seconds), regulation instruction (7 seconds), image viewing (1 second), and affect rating (2.5 seconds). As illustrated in Figure 1A, the trial structure was adapted from several well-established emotion regulation paradigms (Ochsner et al., 2004; Wager et al., 2008; Pitskel et al., 2011; Belden et al., 2014), which reliably elicit amygdala activation (Silvers et al., 2015).

**Figure 1.**
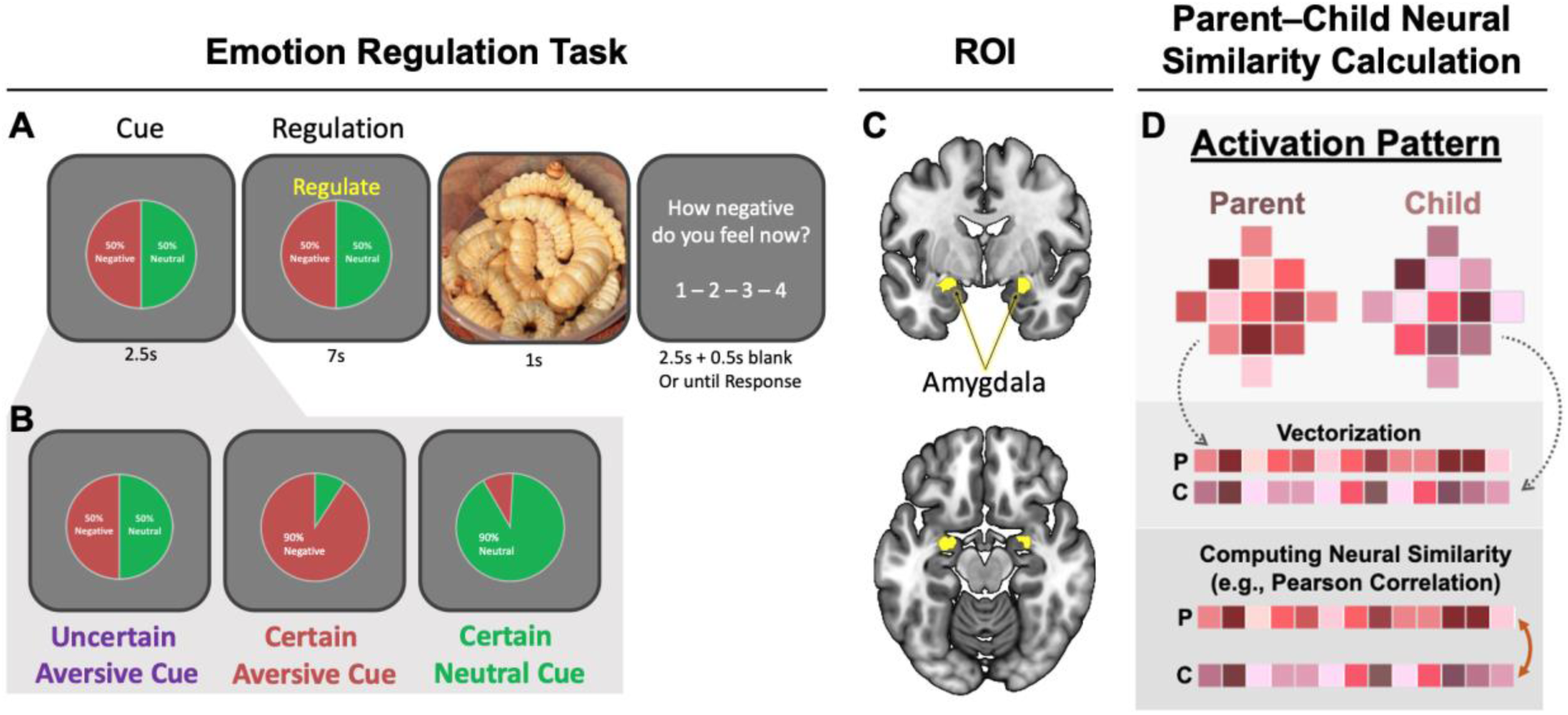
Overview of Experimental Paradigm and Steps of Neural Similarity Calculation (A) Example trial from the Emotion Regulation Task. The upper panel shows the overall trial structure, including the cue, regulation, image presentation, and rating phases. (B) The lower panel displays the three cue types: uncertain aversive, certain aversive, and certain neutral cues, represented by pie charts indicating the probability of viewing a negative or neutral image. (C) The bilateral amygdala was defined as the region of interest (ROI) for neural similarity analyses. (D) Parent–child neural similarity was computed by extracting voxel-wise activation patterns within the amygdala ROI during the task, vectorizing the patterns, and calculating Pearson correlations between parent and youth activation vectors.

The primary aim of the current study was to examine affective processing under uncertainty. To achieve this, the design featured three conditions: (1) an *Uncertainty Aversive Condition*, in which an ambiguous cue was used to elicit uncertainty-related anxiety; (2) a *Certain Aversive Condition*, involving a clearly negative cue; and (3) a *Certain Neutral Condition*. Conditions (2) and (3) served as control conditions to determine whether the observed effects were specific to uncertainty, rather than reflecting general affective responses.

At a cue, participants were signaled the likelihood of viewing a certain aversive image (e.g., wound, snake, or car accident), a certain neutral image (e.g., objects like a chair or cup, or neutral scenes), or an ambiguous cue indicating that the upcoming image could be either negative or neutral (uncertain aversive condition) (Figure 1B).

A regulation phase followed the cue phase to isolate the effects of explicit emotion regulation from early bottom-up arousal elicited by the cue. All conditions were evenly split between “Regulate” and “Maintain” instructions. If presented with “Regulate,” participants were prompted to use any emotion regulation strategies, except for closing their eyes or looking away from the screen, to manage their emotional response. If presented with “Maintain,” they were instructed to keep their emotional state as it was.

Each session included 8 trials per condition (24 trials total) and lasted approximately 11 to 12 minutes. Participants completed three sessions in total. Additional task details, including stimulus timing and post-scan ratings, are available in the Supplementary Materials Section A.

## MRI data acquisition and analysis

### Acquisition

MRI data were collected using a Siemens 3T PRISMA MRI scanner equipped with a 64-channel matrix head coil located at the Fralin Biomedical Research Institute at Virginia Tech Carilion. High-resolution T1 and T2 anatomical images were acquired with an isotropic voxel size of 1 mm and a field of view (FOV) of 256 mm. T1 was scanned with TR = 2.5s, TE = 2.06ms, and flip angle (FA) = 8°; while T2 was scanned with TR = 3.2s, TE = 563ms, and FA = 120°. These images were used for tissue segmentation and normalization. Functional images were acquired for the Emotion Regulation Task (354 volumes), using a gradient-echo echo-planar T2*-weighted imaging sequence with a voxel size of 2.5 x 2.5 mm, 37 interleaved 3.0 mm slices with a 0.3mm gap, FOV = 92 mm, TR = 2s, TE = 25ms, FA = 90°.

### fMRI Preprocessing

The FMRIB Software Library (FSL) (Smith et al., 2004), ICA-AROMA toolbox (Pruim et al., 2015), and ANTs library (Avants et al., 2009) were used for preprocessing. The three task sessions underwent preprocessing, including removal of the first two volumes, motion correction, grand-mean intensity normalization, registration to a standard MNI 2-mm brain template, 5-mm smoothing, ICA denoising, and application of a high-pass filter (0.0078 Hz).

### General Linear Model

SPM12 was utilized to perform data analysis. A general linear model (GLM) with three task sessions was computed using the preprocessed images. We modeled the timing of the cue phase (uncertain aversive cue, certain aversive cue, and certain neutral cue) and the timing of the regulation phase (regulate or maintain). The remaining scenes were included as non-interest events (i.e., nuisance regressors), including the timing of showing negative or neutral pictures and rating, as well as motion regressors comprising six motion parameters and motion outlier time points that exceeded 1.5 times the interquartile range (IQR) from the 75th percentile.

#### Estimation of parent–child neural activation pattern similarity with amygdala seed

The primary interest of the current study was how the amygdala neural similarity between parent and youth associated with youth’s emotional adjustment and parent–youth relationships. To this end, we first defined an amygdala seed (Figure 1B) using 50% thresholded amygdala mask from the Harvard-Oxford subcortical structural atlases (Desikan et al., 2006) and the Neurosynth database (Yarkoni et al., 2011). Given that the Harvard-Oxford Atlas is structurally defined, only the voxels that are also functionally associated with “emotional response” at a *Z* = 3.4 threshold level in the Neurosynth database (included 97 studies, association inference) were included (https://neurosynth.org/analyses/terms/emotional%20responses/). This combined approach represents the amygdala seed associated with emotional response with voxel size *k* = 143. For each participant, the voxel-wise pattern of amygdala activation was extracted and vectorized for the following conditions: *phase of three different cue types* and *phases of emotion regulation* for each aversive cue. The similarity in activation patterns between parents and their children for each phase was calculated using Pearson correlation (*r*) between the parent vector and the child vector (Dimsdale-Zucker and Ranganath, 2018; Lee et al., 2018). A similarity value of 1 indicates identical patterns, while −1 indicates dissimilar patterns. Fisher’s *z* transformation (i.e., *r*-to-*z* transformation) was then performed for further analyses.

## Psychological measures

Youth completed a battery of self-report measures to assess their level of emotional adaptation (i.e., depression, anxiety, and ego resilience) and family relationship qualities (i.e., parent–child conflict, parent–child cohesion, and family identity). Parents were asked to provide additional demographic information, including age, biological sex, and the race and ethnicity of both themselves and their child. This information was used for descriptive purposes to characterize the sample and were included in sensitivity analysis as confounding covariates in the regression models to account for their potential influence.

### Youth Emotional Adaptation

A composite emotional adaptation score was calculated by standardizing three emotional outcomes (depression, anxiety, and ego resilience) into *z*-scores, reversing the depression and anxiety scores, and summing the three *z*-scores (Andrade, 2021). Higher scores indicate better emotional adaptation. The main analyses were based on this composite emotional adaptation score.

### Youth Depression

Youth depression was assessed using the Center for Epidemiologic Studies Depression Scale for Children (Radloff, 1977; Roberts et al., 1990). Youth reported on 20 items, rating how often they experienced each feeling over the past week (e.g., “I was bothered by things that usually don’t bother me,” “I felt like I couldn’t pay attention to what I was doing,” and “I felt down and unhappy.”) on a 4-point Likert scale ranging from 0 (rarely or none of the time [less than 1 day]) to 3 (most or all of the time [5-7 days]). The item scores were averaged, and higher mean scores reflected greater depression (α = 0.83).

### Youth Anxiety

Youth anxiety was evaluated using the Revised Children’s Manifest Anxiety Scale (Reynolds and Richmond, 2011). Youth self-responded to 25 items, rating how often they experienced each feeling over the past week (e.g., “I worried a lot,” “I got nervous when things did not go the right way,” and “I worried about what was going to happen to me.”) on a 5-point Likert scale ranging from 0 (never) to 4 (very often). The item scores were averaged, and higher mean scores reflected greater anxiety (α = 0.92).

### Ego Resilience

Youth ego resilience was assessed using the 6-item Brief Resilience Scale (Smith et al., 2008). Youth self-rated how often they experienced each feeling over the past week for each item (e.g., “I have a hard time making it through stressful events,” “I tend to bounce back quickly after hard times,” and “It is hard for me to snap back when something bad happens.”) on a 5-point Likert scale from 1 (strongly disagree) to 5 (strongly agree). The item scores were averaged, with higher mean scores indicating greater ego resilience and ability to bounce back from negative affect (α = 0.85).

### Youth-Perceived Family Relationships

A composite family relationships score was calculated by standardizing three family relationship qualities (parent–youth conflict, parent–youth cohesion, and family identity) into *z*-scores, reversing the conflict scores, and summing the three *z*-scores (Andrade, 2021). Higher scores indicate better youth-perceived family relationships. The main analyses were based on this composite family relationships score.

### Parent-Youth Conflict

Youth-reported relationship conflict was assessed using the scale developed by Ruiz et al. (1998), which measures the frequency of arguments and fights between parents and children (Ruiz et al., 1998). Participants were instructed to reflect on their experiences specifically with the parent who participated in the scan with them. Participants responded to 10 items, rating how often they experienced each situation (e.g., “You and your parents disagreed with each other,” “You and your parents ignored each other,” and “You and your parents yelled or raised your voices at each other”) over the past month on a 5-point Likert scale ranging from 1 (almost never) to 5 (almost always). The item scores were averaged, with higher mean scores reflecting greater parent-youth conflict (α = 0.88).

### Parent-Youth Cohesion

Youth-reported cohesion was assessed using the Cohesion subscale of the Family Adaptation and Cohesion Evaluation Scales II inventory (Olson et al., 1979), which measures the relationship closeness between youth and their parents. Specifically, youths rated how strongly they agreed or disagreed with each statement (e.g., “My mother/father and I are supportive of each other during difficult times,” “My mother/father and I do things together,” and “My mother/father and I like to spend our free time with each other”) based on their experiences with the parent who participated in the scan with them. Responses were recorded on a 5-point Likert scale, ranging from 1 (almost never) to 5 (almost always). The item scores were averaged, with higher mean scores reflecting greater parent–youth cohesion (α = 0.81).

### Youth Family Identity

Youth-reported family identity was assessed using an 8-item scale measuring the extent to which an individual’s sense of self is connected to their family values (Tyler and Degoey, 1995). Specifically, participants rated how strongly they agreed or disagreed with each statement (e.g., “My family is important to the way I think of myself as a person,” “I feel that my parents respect the work I do,” and “I feel like a valued member of my family”) based on their experiences with the parent who participated in the scan with them.

Responses were recorded on a 5-point Likert scale, ranging from 1 (strongly disagree) to 5 (strongly agree). The item scores were averaged, with higher mean scores reflecting a stronger sense of family identity (α = 0.86).

## Analytic Plan

### Examining Hypotheses

All analyses for examining hypotheses were carried out using JASP (JASP Team, 2024). To test the hypotheses, the regression model with 5,000 bootstrapping resampling at a 95% confidence interval (*p* < .05) was applied. Bootstrapping is a resampling technique that helps accommodate small sample sizes by generating empirical confidence intervals without relying on normality assumptions or large-sample theory (Efron and Tibshirani, 1993). A non-zero value within the confidence interval suggests evidence of a significant effect. When each component of the composite scores was tested separately, false discovery rate (FDR) correction using the Benjamini–Hochberg procedure (Benjamini and Hochberg, 1995) was applied with a threshold of Q = .05. In these cases, bias-corrected and accelerated (BCa) *p*-values derived from the bootstrap distribution (Efron, 1987) were used. The BCa *p*-values were ranked in ascending order and compared against their corresponding FDR-adjusted critical values to determine significance. This procedure controls the expected proportion of false positives among the rejected hypotheses while preserving statistical power in the presence of multiple comparisons.

#### Prefrontal Recruitment

A generalized psychophysiological interaction (gPPI) analysis was conducted for the youth participants. The amygdala-seed-based whole-brain analysis with parent–child amygdala similarity as a regressor was performed. A voxel-level threshold of uncorrected *p* < .005 and a familywise error (FWE) cluster-level correction at *p* < .05 at the whole-brain level were set. Significant clusters were considered part of the amygdala–prefrontal inhibitory system only if they overlapped with regions defined by both the Automated Anatomical Labeling (AAL) atlas (Rolls et al., 2020) and Brodmann Areas. Specifically, eligible clusters had to fall within the following AAL regions: Superior Frontal Gyrus, Middle Frontal Gyrus, Inferior Frontal Gyrus, or Medial Frontal Gyrus, and simultaneously overlap with Brodmann Areas 9, 10, 11, 44, 45, 46, or 47.

### Sensitivity Analysis

To ensure the results specifically reflected parent–child neural similarity during uncertainty-induced affective processing, additional analyses examined parent–child neural similarity during the certain aversive cue, the neutral cue, and between youths and non-parent adults. To address potential confounds, all significant findings were retested while controlling for youth age, biological sex (female = 1, male = 0), and whether the parent–child pair was of the same sex (1) or of different sexes (0).

## Results

Descriptive statistics and correlations among demographic variables and study measures are reported in Supplementary Materials Section B (Table S1).

### Parent–Child Dyadic Neural Similarity and Youth’s Emotional Adaptation

The first set of analyses examined whether parent–child amygdala activation similarity during uncertainty-induced affective processing was associated with youth emotional adaptation. The cue phase and regulation phase were analyzed separately to isolate the effects of explicit emotion regulation from early bottom-up arousal elicited by the cue.

### Uncertain Aversive Cue

Regression analysis showed that parent–child similarity in amygdala activation during uncertainty-induced affective processing was positively associated with youths’ emotional adaptation (B = 3.550, SE = 1.212, 95% CI = [1.490, 6.464], Figure 2A). The result remained significant when accounting for the effects of youth age, biological sex, and whether the parent– child pair was of the same sex (B = 3.314, SE = 1.502, 95% CI = [0.876, 7.004]). Furthermore, each component of the emotional adaptation score (i.e., depression, anxiety, and ego resilience) was examined individually, and all results remained significant after FDR correction: greater neural similarity was associated with lower depression and anxiety, and higher ego resilience. Detailed statistical results for each emotional profile are provided in Supplementary Materials Section C-1.

**Figure 2.**
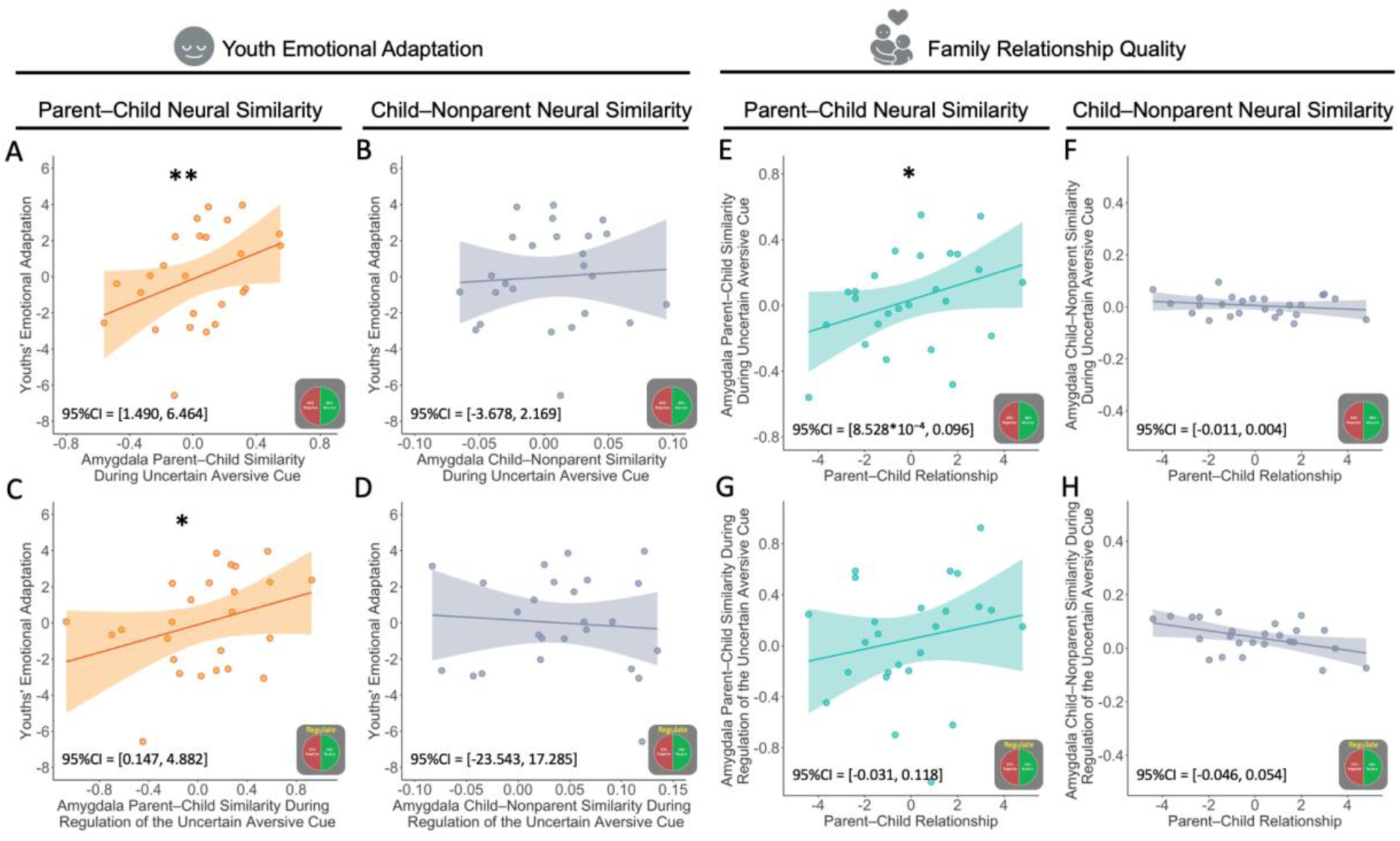
Associations Between Parent–Child Amygdala Activation Neural Similarity, Youth Emotional Adaptation, and Family Relationship Quality. (A-D) Scatter plots show the analyses examining the association between amygdala neural similarity and youths’ emotional adaptation. (E-H) Associations between youth-perceived family relationship quality and dyadic amygdala similarity. Shaded regions represent 95% confidence intervals (CI). \**p* < .05, \*\**p* < .01 (bias-corrected and accelerated *p*-values, estimated from 5,000 bootstrap resamples).

### Sensitivity Test

Results of the sensitivity analyses showed that parent–child similarity in amygdala activation during the certain aversive cue (B = 2.631, SE = 1.548, 95% CI = [0.231, 6.598]) was significantly associated with youths’ emotional adaptation. However, the result was no longer significant when accounting for youth age, biological sex, and whether the parent– child pair was of the same sex (B = 2.390, SE = 1.883, 95% CI = [-0.635, 7.095]). No significant associations were found for the neutral cue (B = −0.273, SE = 1.418, 95% CI = [-3.678, 2.169]) or between youths and non-parent adults (B = 4.155, SE = 11.555, 95% CI = [-16.492, 29.553], Figure 2B). These results suggest that the association between parent–child dyadic neural similarity during uncertainty-induced affective processing and youths’ emotional adaptation is more robust and reliable.

### Regulation of the Uncertain Aversive Cue

Regression analysis showed that parent–child similarity in amygdala activation during the regulation of uncertainty-induced affective processing was positively associated with youths’ emotional adaptation (B = 1.890, SE = 1.165, 95% CI = [0.147, 4.882], Figure 2C). When accounting for the effects of youth age, biological sex, and whether the parent–child pair was of the same sex, marginal significance was observed (B = 1.846, SE = 1.210, 95% CI = [-0.015, 4.805]). Furthermore, each component of the emotional adaptation score (i.e., depression, anxiety, and ego resilience) was examined individually. After applying FDR correction, significant associations were found for depression and anxiety only but not for ego resilience. Detailed statistical results for each emotional profile are provided in Supplementary Materials Section C-2.

### Sensitivity Test

Results of the sensitivity analyses showed that parent–child similarity in amygdala activation during the regulation of the certain aversive cue was significantly associated with youths’ emotional adaptation (B = 2.170, SE = 0.863, 95% CI = [0.616, 3.986]). This association remained significant when controlling for youth age, sex, and pair type (B = 2.217, SE = 1.009, 95% CI = [0.386, 4.395]). No significant association was found between youths and non-parent adults (B = −3.581, SE = 10.562, 95% CI = [-23.543, 17.285], Figure 2D). This suggests that the association between parent–child neural similarity during the regulation of uncertain aversive cues and adolescents’ emotional adjustment is not specific, as a similar effect was also observed during the regulation of certain aversive cues.

## Family Relationships and Parent–Child Dyadic Neural Similarity

The second set of analyses examined whether youth-perceived family relationships were associated with parent–child amygdala activation similarity during uncertainty-induced affective processing. Consistent with the previous set of analyses, the cue phase and regulation phase were analyzed separately to isolate the effects of explicit emotion regulation from early bottom-up arousal elicited by the cue.

### Uncertain Aversive Cue

Regression analysis showed that youth-perceived family relationships were positively associated with parent–child similarity in amygdala activation during uncertainty-induced affective processing (B = 0.043, SE = 0.024, 95% CI = [8.528*10⁻⁴, 0.096], Figure 2E). This association remained significant after controlling for youth age, biological sex, and parent–child sex pair type (B = 0.050, SE = 0.028, 95% CI = [0.002, 0.111]). Furthermore, each component of the family relationship quality score (conflict, cohesion, and identity) was examined individually to assess their specific contributions. Results showed that only parent–child conflict was significantly associated with amygdala similarity after FDR correction, such that lower conflict was linked to greater neural similarity. Detailed statistical results for each relationship component are provided in Supplementary Materials Section C-3.

### Sensitivity Test

Sensitivity tests were conducted to examine parent–child neural similarity during the certain aversive cue, the neutral cue, and between youths and non-parent adults. Results showed that youth-perceived family relationships were not significantly associated with neural similarity during the certain aversive cue (B = 0.001, SE = 0.026, 95% CI = [-0.039, 0.067]), the neutral cue (B = −0.002, SE = 0.025, 95% CI = [-0.046, 0.054]), or between youths and non-parent adults (B = −0.004, SE = 0.004, 95% CI = [-0.011, 0.004], Figure 2F). These results suggest that the observed associations are specific to parent–child dyads during uncertainty-induced affective processing.

### Regulation of the Uncertain Aversive Cue

Regression analysis showed that youth-perceived family relationships were not significantly associated with parent–child similarity in amygdala activation during the regulation of uncertainty-induced affective processing (B = 0.038, SE = 0.037, 95% CI = [-0.031, 0.118], Figure 2G).

## Mediating Role of Parent–Child Neural Similarity

### Uncertain Aversive Cue

Given that previous studies suggest family relationships as an independent variable and parent–child neural similarity as a mediator (Lee et al., 2018), Model 1 in this mediation analysis applied the same setup. Specifically, Model 1 (Figure 3A) was conducted to examine whether parent–child neural similarity during uncertainty-induced affective processing mediated the association between youth-perceived family relationships and their emotional adaptation. A significant indirect effect was found (B = 0.111, SE = 0.096, 95% CI = [0.004, 0.354]); while the direct effect of youth-perceived family relationships on emotional adaptation was not significant (B = 0.360, SE = 0.213, 95% CI = [-0.145, 0.952]), suggesting that parent–child neural similarity fully mediated this association. The indirect effect accounted for 23.57% of the total effect (B = 0.471, SE = 0.206, 95% CI = [0.025, 1.022]).

**Figure 3.**
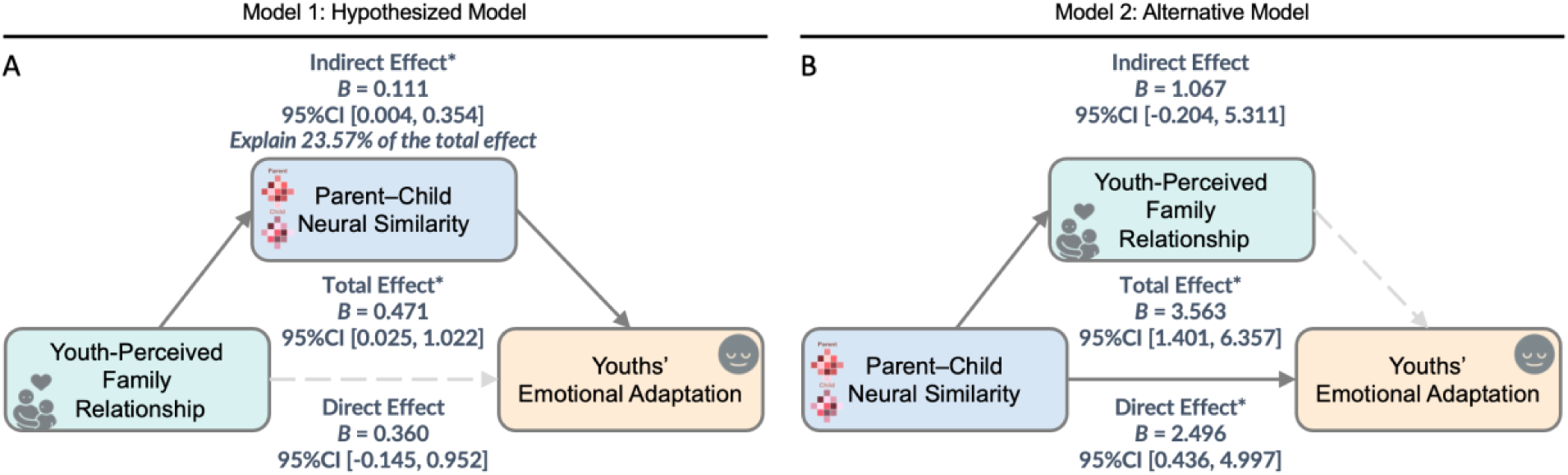
Mediation Models Examining the Role of Parent–Child Neural Similarity in Amygdala and Youth-Perceived Family Relationships in Predicting Youths’ Emotional Adaptation. (A) Hypothesized mediation model: Youth-perceived family relationship quality is positively associated with parent–child neural similarity in the amygdala during the uncertain aversive cue phase, which in turn is associated with better youth emotional adaptation. The indirect effect was significant, accounting for 23.57% of the total effect. (B) Alternative model: Parent–child neural similarity predicts family relationship quality, which then predicts emotional adaptation. The indirect path was not significant. * denotes significant effects (95% Confidence intervals does not include zero). Shaded paths indicate non-significant effects.

To further explore the direction of the mediation pathway, a similar mediation analysis was conducted with parent–child neural similarity as the independent variable and youth-perceived family relationships as the mediator (Model 2, Figure 3B). However, the indirect effect through youth-perceived family relationships was not significant (B = 1.067, SE = 0.837, 95% CI = [-0.204, 5.311]). These findings suggest that youth-perceived family relationships did not significantly mediate the association between parent–child neural similarity during uncertainty-induced affective processing and youths’ emotional adaptation.

A supplementary analysis was conducted using parent–child conflict as a predictor, as it was the only relationship quality that showed a robust association with parent–child neural similarity during uncertainty-induced affective processing. Consistent with the main mediation model, the analysis revealed a significant indirect effect (B = −0.493, SE = 0.395, 95% CI = [-1.534, −0.060]), suggesting that parent–child neural similarity fully mediated the association between parent–child conflict and youth emotional adaptation (Figure S1A). Detailed statistical results for each relationship component are provided in Supplementary Materials Section C-4.

### Regulation of the Uncertain Aversive Cue

Model 1 tested whether parent–child neural similarity during the regulation of the uncertain aversive cue mediated the link between youth-perceived parent–child relationships and emotional adaptation. The indirect effect was not significant (B = 0.420, SE = 0.460, 95% CI = [-0.256, 2.385]). Model 2 examined another mediation pathway, with neural similarity as the independent variable and perceived family relationships as the mediator. This model was also not significant in indirect effect (B = 0.059, SE = 0.071, 95% CI = [-0.032, 0.304]), suggesting no evidence of mediation in either direction.

## Influence of Amygdala Parent–Child Neural Similarity on Youth’s Prefrontal Recruitment

An amygdala-seed-based whole-brain gPPI connectivity analysis was conducted using parent–child amygdala activation pattern similarity during the uncertain aversive cue as a regressor. This analysis aimed to examine whether neural similarity is associated with youth prefrontal engagement during uncertain situations, addressing a mechanistic question of how shared affective processing between parent and child may support regulatory functioning in youth. The contrast of Regulation versus Cue under the uncertainty condition (i.e., Regulation of Uncertain Cue > Uncertain Cue) was used to isolate regulation-related neural mechanisms.

### Regressor: Neural Similarity During Uncertain Aversive Cue

The result revealed that parent–child amygdala similarity was negatively associated with youth amygdala connectivity to several frontal and motor regions (Figure S2). Specifically, significant clusters were observed in the right dlPFC (Table 1, Cluster 1), the somatomotor and sensory cortex (Table 1, Cluster 2&4), and the supplementary motor area (SMA; Table 1, Cluster 3). The coordinate information is presented in Table 1.

**Table 1.**
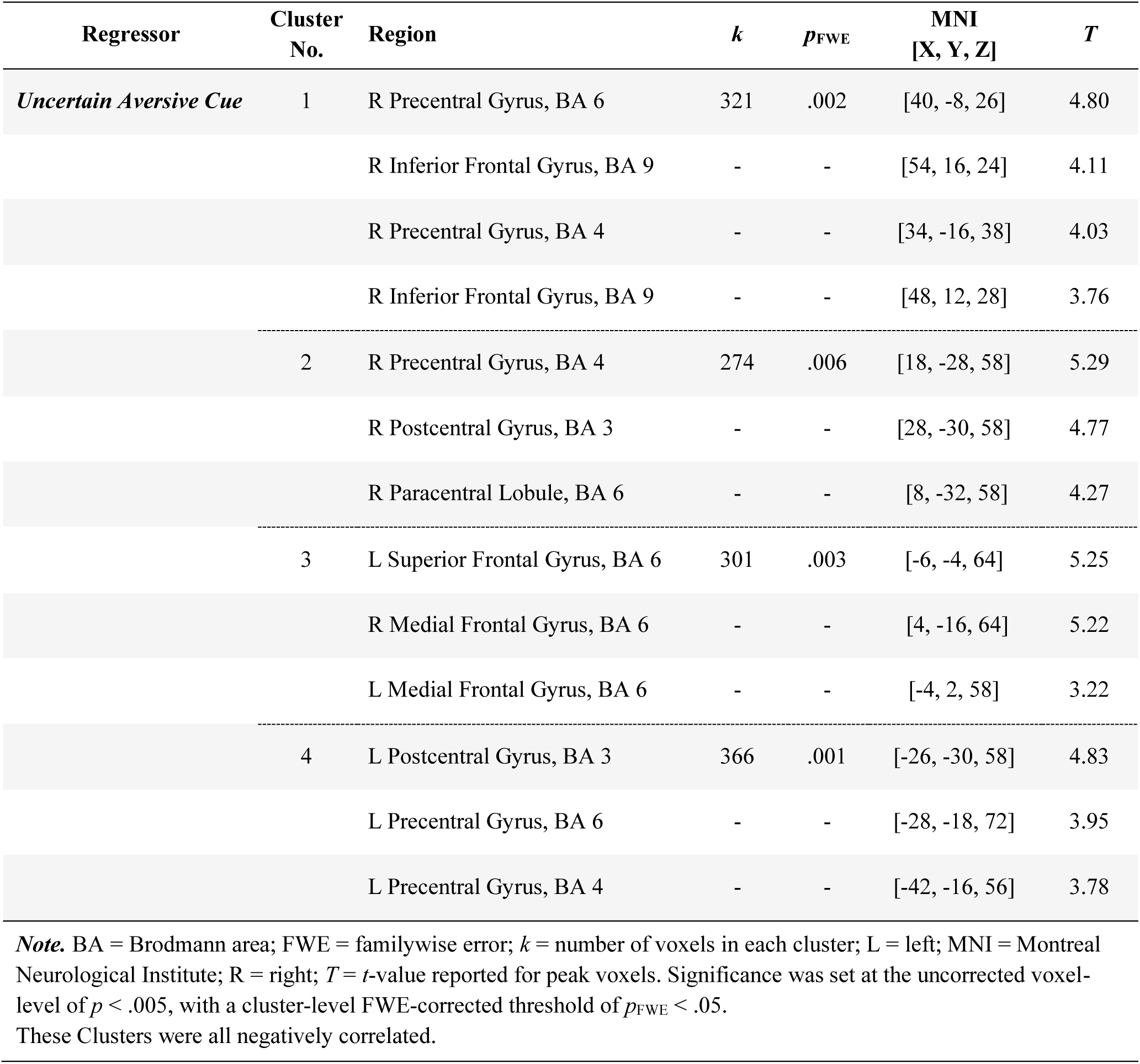
Significant Clusters Identified in the Amygdala-Seed-Based Whole-Brain gPPI Connectivity Analysis with the Contrast of Regulation of Uncertain Aversive Cue Minus Uncertain Aversive Cue

Given that amygdala-dlPFC connectivity (Figure 4A; Figure S2) was identified using the contrast of regulation minus cue during the uncertainty condition, we extracted connectivity values separately for the uncertain aversive cue phase and the regulation phase to determine which phase was driving the effect. These phase-specific connectivity values were then used to predict youths’ emotional adaptation. The results showed that greater amygdala-dlPFC connectivity during the cue phase (B = 3.300, SE = 1.373, 95% CI = [1.008, 6.478], Figure 4B), but not during the regulation phase (B = −1.775, SE = 4.953, 95% CI = [-13.994, 6.179], Figure 4C), was associated with better emotional adaptation. The association between cue-phase connectivity and emotional adaptation remained significant after FDR correction, and when controlling for youth age, biological sex, and pair type (B = 3.432, SE = 1.694, 95% CI = [0.228, 6.948]).

**Figure 4.**
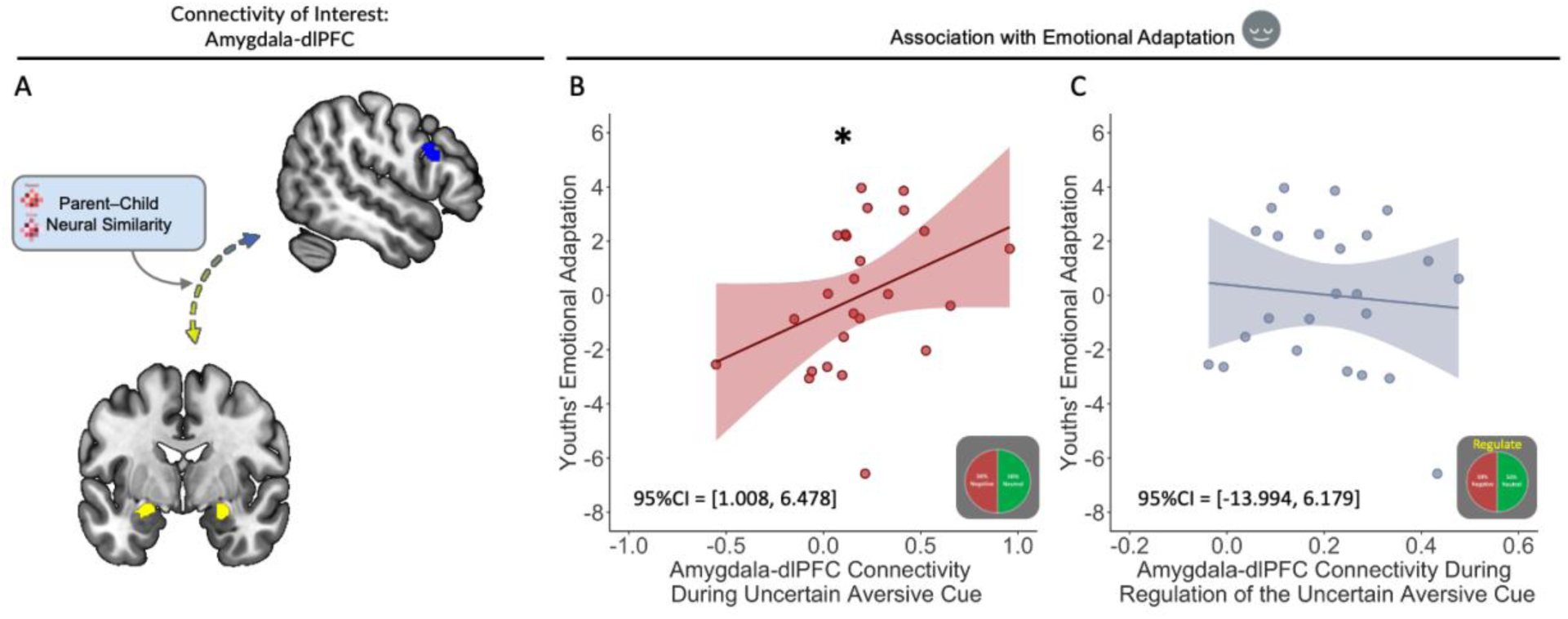
Amygdala-dlPFC Connectivity and Youths’ Emotional Adaptation (A) The connectivity of interest between the amygdala (seed region, in yellow) and the dorsolateral prefrontal cortex (dlPFC, in blue) is illustrated. (B) Greater amygdala-dlPFC connectivity during the uncertain aversive cue phase was significantly associated with higher youths’ emotional adaptation. (C) No significant association was found between amygdala-dlPFC connectivity during the regulation phase of the uncertain cue and youths’ emotional adaptation. Shaded regions represent 95% confidence intervals (95%CI). \**p* < .05 (bias-corrected and accelerated *p*-values, estimated from 5,000 bootstrap resamples).

### Regressor: Neural Similarity During Regulation of Uncertain Aversive Cue

A gPPI analysis using parent–child amygdala similarity during the regulation phase as a regressor revealed negative associations with amygdala connectivity to motor-related regions, based on the contrast comparing regulation to cue presentation in uncertain aversive conditions (Figure S3). Specifically, significant negative associations were observed in the SMA and midcingulate cortex (Table 2, Cluster 1) and the temporal cortex (Table 2, Cluster 2), with no significant effects found in prefrontal regions. Full coordinate details are presented in Table 2.

**Table 2.** Significant Clusters Identified in the Amygdala-Seed-Based Whole-Brain gPPI Connectivity Analysis with the Contrast of Regulation of Uncertain Aversive Cue Minus Uncertain Aversive Cue

| Regressor | Cluster No. | Region | *k* | *p*<sub>FWE</sub> | MNI [X, Y, Z] | *T* |
| --- | --- | --- | --- | --- | --- | --- |
| *Regulation of Uncertain Aversive Cue* | 1 | R Medial Frontal Gyrus, BA 6 | 463 | <.001 | [6, -20, 56] | 5.30 |
|  |  | L Medial Frontal Gyrus, BA 6 | - | - | [-2, 0, 58] | 4.87 |
|  |  | R Cingulate Gyrus, BA24 | - | - | [10, 0, 42] | 3.95 |
|  | 2 | L Superior Temporal Gyrus, BA 22 | 281 | .005 | [-50, 0, -4] | 5.01 |
|  |  | L Middle Temporal Gyrus, BA 21 | - | - | [-60, -6, -10] | 4.48 |
|  |  | L Inferior Temporal Gyrus, BA 21 | - | - | [-58, -12, -16] | 3.70 |

**Note.** BA = Brodmann area; FWE = familywise error; *k* = number of voxels in each cluster; L = left; MNI = Montreal Neurological Institute; R = right; *T* = *t*-value reported for peak voxels. Significance was set at the uncorrected voxel-level of $p < .005$ , with a cluster-level FWE-corrected threshold of $p_{\text{FWE}} < .05$ . These Clusters were all negatively correlated.

## Discussion

Adolescence is a period marked by heightened uncertainty, which can increase vulnerability in youth (Brown, 2004; Dugas et al., 2004; Byrne et al., 2007). The current study aimed to investigate whether neural similarity between youth and their parents serves as a protective factor for emotional adaptation, and to explore the role of the family relationship in shaping this neural alignment. As hypothesized, the results indicated that parents and youths from more positive family relationships, particularly those perceived lower parent–child conflict, showed greater neural similarity in the amygdala when facing uncertainty. In turn, higher parent–child amygdala similarity during these uncertainty cues was associated with better youth’s emotional adaptation, including lower anxiety and depression as well as higher resilience. In addition, the current study incorporated the dual-systems model of adolescent brain development to explore a potential neural mechanism linking affective similarity and regulatory function. Specifically, the functional connectivity results demonstrated that parent– child amygdala similarity was associated with youth’s greater dlPFC engagement during uncertainty, suggesting enhanced top-down regulation.

The finding of higher parent–child neural similarity having a protective effect aligns with current literature on parent–child neural similarity, which suggests such similarity is associated with reduced stress and internalizing symptoms (Lee et al., 2018; Quiñones-Camacho et al., 2022), better emotional functioning (Lee et al., 2017a, 2017c), greater resilience (Zhou et al., 2023), and protection from risky behaviors (Kim-Spoon et al., 2024). For example, one recent fMRI study found that when parents and children exhibited similar connectivity in emotion-related networks while watching emotional films, youth reported less anxiety and greater ego-resilience, but only in families with high cohesion (Zhou et al., 2023). However, prior studies have not tested the full process from family relationship quality to parent–child neural similarity and, in turn, to youth’s emotional adaptation. The current study extends this literature by providing evidence of a mediating pathway. Despite its cross-sectional design, the findings suggest that shared neural responses mediate the association between supportive family environments and adolescent emotional health. In other words, having a parent “fire together” neurologically during stressful uncertainty may help youths manage emotional challenges more effectively. Although we examined an alternative model suggesting that family relationships may shape emotional adaptation through neural similarity, longitudinal research is needed to establish the temporal sequence of these associations and to confirm whether positive family relationships can cultivate parent–child neural similarity and, in turn, enhance youth emotional adaptation.

This “firing-together” amygdala activation was also associated with prefrontal recruitment in the early emotional arousal stage (i.e., cue phase). Higher parent–child amygdala similarity was linked to youth’s increased amygdala-dlPFC connectivity specifically during the uncertain cue phase but not the explicit effortful regulation phase. Moreover, youths who demonstrated stronger amygdala-dlPFC coupling during this early emotional arousal stage exhibited better emotional outcomes. These findings echo the social developmental theory of co-regulation, which posits that supportive caregivers actively modulate a child’s emotional and physiological arousal, gradually scaffolding the child’s capacity for self-regulation through sensitive interactions (Feldman, 2012; Reindl et al., 2018). Simultaneously, these results align with social buffering theory, highlighting that the synchronous emotional response of a trusted caregiver can reduce physiological and neural stress responses, especially in uncertain or threatening contexts (Hostinar et al., 2014; Gunnar and Hostinar, 2015). Our findings extend both theories by proposing a possible neural mechanism: amygdala parent–child neural similarity in emotional arousal may facilitate implicit emotion regulation (Tottenham, 2015). Specifically, “firing together” in emotionally salient moments may automatically recruit prefrontal resources (i.e., dlPFC), reducing the regulatory burden without requiring consciously effortful control (Gyurak et al., 2011; Colich et al., 2017). The current results suggest that shared neural activation patterns during initial emotional arousal stages may represent a fundamental neurodevelopmental mechanism through which caregivers foster effective implicit emotion regulation in youths.

Theoretically, the current findings lend support to the dual-systems model of adolescent development by illustrating the interplay between subcortical emotional reactivity, regulatory control, and the socioenvironmental context. According to the dual-systems model (Casey et al., 2008; Steinberg et al., 2008), adolescence is marked by heightened reactivity of affective brain systems (e.g., the amygdala and reward centers) relative to still-maturing prefrontal control systems. Our findings suggest that positive family relationship and parent–child interactions may help mitigate this imbalance. Specifically, when a youth’s surging amygdala response is aligned with their parent’s response, it might indicate an external regulatory influence that keeps their emotional reactions appropriately calibrated.

Greater parent–child interaction offers more opportunities for youths to learn from and then align with their parents. However, the quality of these interactions, whether positive or negative, and how youths perceive themselves within the family context may lead to different outcomes (Su et al., 2024). Despite growing interest in the link between parent–child neural similarity and youth emotional outcomes, few empirical studies have investigated whether distinct dimensions of family relationship exert comparable effects on neural alignment and youth well-being. Existing research has primarily focused on positive family factors, such as emotional support and time spent together (Zhou et al., 2023), and the personal importance of family values (Lee et al., 2018), while relatively less is known about the influence of negative dynamics such as conflict. The current study addressed this gap by examining three dimensions of family relationship quality (i.e., parent–child conflict, parent–child cohesion, and family identity) to provide a more comprehensive picture of how relational context shapes shared neural responses. Among these, only parent–child conflict showed a robust association with amygdala similarity. This finding aligns with recent research in late childhood, which demonstrated that negative, but not positive, family emotional climate was associated with altered parent–child hippocampal connectivity and increased risk for child psychopathology (Su et al., 2024). Extending this work into adolescence, the present study found that higher levels of parent–child conflict predicted reduced neural similarity in the amygdala during emotionally uncertain situations and increased risk for youth psychopathology. These findings highlight the significance of relational quality in shaping shared neural responses and their implications for youth emotional adaptation, suggesting that future studies should consider multiple dimensions of family relationship quality to better understand their distinct contributions to neural and emotional development.

Although resting-state and passive tasks are commonly used in developmental neuroimaging due to their feasibility in the MRI scanner (Lee et al., 2017b; Qu et al., 2023; Zhou et al., 2023; Su et al., 2024), the current findings highlight the value of context-sensitive tasks in addressing mechanistic questions. Such tasks may be better suited to capture dynamic neural processes, particularly given that real-world experiences are inherently context-rich. Supporting this view, the present study identified several context-specific effects. Associations between parent–child neural similarity, family relationship, and youth’s emotional adaptation were observed specifically during uncertain aversive conditions, but not under certain aversive or neutral conditions, and during the early arousal stage (i.e., cue phase), rather than during the explicit emotion regulation phase. Furthermore, analyses of neural connectivity revealed that greater parent–child amygdala similarity was associated with increased engagement of the dlPFC during the uncertain cue phase. These effects were not observed during the later regulation phase. Together, these context-specific findings underscore the value of task-based fMRI designs in parent–child neural similarity research, demonstrating how such paradigms can capture distinct neural dynamics that may be overlooked in more static or passive approaches.

While this study offers important insights, several limitations should be acknowledged. First, the cross-sectional and correlational design limits causal inference. Although mediation analyses supported a model in which family context influences neural similarity, which in turn supports youth emotional adaptation, longitudinal studies are needed to test these developmental pathways over time. Therefore, any implication that cultivating specific family relationship qualities can alter neural similarity should be interpreted with caution until confirmed by future longitudinal research. Second, some neural similarity may reflect shared genetics rather than relational attunement. Studies using twin or adoptive samples could help disentangle genetic and environmental contributions. Lastly, the modest sample size limited our ability to examine effects based on parent–child sex pairings or developmental differences across stages of adolescence. Future research should consider larger samples and group youth by age to explore developmental differences.

In conclusion, the current study provides evidence that a positive family relationship, particularly one characterized by low conflict, is associated with parent–youth neural similarity in affective processing regions. This neural alignment may serve as a protective factor for youth mental health. Furthermore, the findings suggest that amygdala similarity supports increased engagement of prefrontal regulatory regions, pointing to a potential neural mechanism underlying implicit emotion regulation during uncertainty. These results help bridge social and neural explanations for emotional resilience, highlighting parent–child attunement as a key neurobiological mechanism supporting emotional adaptation. Together, this research advances our understanding of how shared neural responses within families contribute to youth well-being and provides a foundation for future work aimed at informing family-based interventions.

## Acknowledgments

This work was supported through a research award by the Virginia Tech Institute for Society, Culture and Environment to T.-H. Lee, and 2023 Alliance for Neurodevelopment Research Seed Funding, Virginia Tech, and Government Scholarships to Study Abroad, the Ministry of Education, Taiwan, to Y.-Y. Chen.

