## supplementary materials for "From Family Relationship to Regulation: Parent–Child Amygdala Activation Similarity is Linked to Prefrontal Recruitment and Youth Emotional Adaptation"

**A. Emotion Regulation Task**

At the start of each trial (cue phase), a pie chart was presented in the middle of the screen for 2 to 3 seconds. There were three conditions for the cue phase. In the Neutral Cue condition, participants saw a pie chart with a larger portion in green, indicating they had a high (75% to 95%) chance of seeing a neutral picture later in this trial. In the Negative Cue condition, participants saw a pie chart with a larger portion in red, indicating they had a high (75% to 95%) chance of seeing a negative picture later in this trial. In the Ambiguous Cue condition, which elicited uncertainty-induced stress, participants saw a pie chart with equal red and green portions, indicating they had an equal (45% to 55%) chance of seeing either a negative or neutral picture later. There were 8 trails for each condition, yielding 24 trails in total for each session.

Next, an instruction appeared above the pie chart for a 7-second interval (regulation phase). The trials were equally divided into the Emotion Regulation condition and the Maintain Emotion condition. In the Emotion Regulation condition, participants saw a "Regulate" prompt, asking them to use any strategies to regulate their emotions, except for closing their eyes or looking away from the screen. In the Maintain Emotion condition, participants saw a "Maintain" prompt, asking them to maintain their current emotional state or keep their feelings as they were. Following a jitter of 0.5 to 1.5 seconds, a neutral or negative picture was shown on the screen for one second. Participants were then prompted to rate "How negative do you feel now?" They had 2.5 seconds to rate their feelings on a scale of 1 to 4, with 1 being the least negative and 4 being the most negative. Responses were made using a 4-button box in the MRI scanner. Each session took 11 to 12 minutes. Participants completed three sessions, with different structural scans (e.g., T1 and T2 scans) in between, serving as rest periods.

After the scan, participants were requested to rate the pie charts with the question "How anxious did you feel when you saw the following picture in the scanner?" using a scale of 0 to 10, with 0 indicating not at all anxious and 10 indicating extremely anxious. Additionally, participants were asked to report their emotion regulation strategies by responding to an open-ended question and a multiple-choice question where they can select all that apply.

### B. Descriptive Statistics and Correlations Among Demographic Variables

**Table 1.** *Correlations and Descriptive Statistics [Mean (SD)] for Demographic Variables and Study 1 Measures*

| | **Youth Age [*r*]** | **Youth Sex** | **Pair Type** | **Male-Male** | **Male-Female** | **Female-Male** | **Female-Female** |
|---|---|---|---|---|---|---|---|
| **Family Relationship** | -.239 | F: -0.05 (2.57)<br>M: 0.04 (2.21) | S: -0.20 (2.50)<br>D: 0.13 (2.28) | 0.16<br>(1.61) | 0.46<br>(2.16) | 0.01<br>(2.42) | -0.35<br>(2.90) |
| Conflict | .065 | F: 2.11 (0.82)<br>M: 2.29 (0.73) | S: 2.39 (0.74)<br>D: 2.09 (0.77) | 2.47<br>(0.64) | 1.68<br>(0.70) | 2.24<br>(0.77) | 2.36<br>(0.83) |
| Cohesion | -.635*** | F: 3.56 (0.85)<br>M: 3.76 (0.55) | S: 3.81 (0.77)<br>D: 3.59 (0.64) | 4.13<br>(0.06) | 3.38<br>(0.85) | 3.66<br>(0.58) | 3.67<br>(0.90) |
| Identity | .013 | F: 4.13 (0.65)<br>M: 4.16 (0.69) | S: 4.05 (0.78)<br>D: 4.22 (0.58) | 4.04<br>(0.96) | 4.28<br>(0.36) | 4.20<br>(0.65) | 4.06<br>(0.78) |
| **Emotional Adaptation** | .058 | F: -0.33 (2.98)<br>M: 0.26 (2.43) | S: -0.56 (3.18)<br>D: 0.37 (2.25) | 0.40<br>(3.40) | 0.80<br>(2.35) | 0.22<br>(2.31) | -0.97<br>(3.27) |
| Depression | .053 | F: 0.86 (0.48)<br>M: 0.68 (0.37) | S: 0.85 (0.51)<br>D: 0.70 (0.35) | 0.50<br>(0.39) | 0.61<br>(0.37) | 0.73<br>(0.36) | 1.00<br>(0.50) |
| Anxiety | -.076 | F: 1.50 (0.77)<br>M: 1.32 (0.64) | S: 1.52 (0.90)<br>D: 1.32 (0.52) | 1.25<br>(1.11) | 1.27<br>(0.59) | 1.34<br>(0.53) | 1.63<br>(0.87) |
| Ego Resilience | .129 | F: 3.18 (0.79)<br>M: 3.10 (0.78) | S: 3.01 (0.86)<br>D: 3.22 (0.72) | 2.80<br>(0.78) | 3.34<br>(0.53) | 3.18<br>(0.80) | 3.09<br>(0.94) |
| Amygdala Similarity (Uncertain Aversive Cue) | .191 | F: -0.004 (0.29)<br>M: 0.065 (0.28) | S: -0.022 (0.24)<br>D: 0.072 (0.31) | 0.081<br>(0.20) | 0.105<br>(0.36) | 0.060<br>(0.31) | -0.066<br>(0.26) |
| Amygdala Similarity (Regulation of Uncertainty) | -.130 | F: 0.079 (0.44)<br>M: 0.031 (0.49) | S: -0.006 (0.39)<br>D: 0.091 (0.51) | 0.069<br>(0.43) | 0.284<br>(0.48) | 0.020<br>(0.52) | -0.039<br>(0.40) |
| Amygdala-Prefrontal Connectivity Strength (Uncertain Aversive Cue) | .259 | F: 0.187 (0.23)<br>M: 0.200 (0.34) | S: 0.183 (0.20)<br>D: 0.202 (0.35) | 0.156<br>(0.199) | 0.175<br>(0.28) | 0.212<br>(0.38) | 0.194<br>(0.21) |
| Amygdala-Prefrontal Connectivity Strength (Regulation of Uncertainty) | -.103 | F: 0.318 (0.13)<br>M: 0.217 (0.43) | S: 0.378 (0.43)<br>D: 0.184 (0.14) | 0.211<br>(0.081) | 0.090<br>(0.06) | 0.219<br>(0.14) | 0.449<br>(0.51) |

***Note.*** **Pair Type:** Indicates whether the parent–child pair was of the same sex or different sexes. **F:** Female ($n$ = 11). **M:** Male ($n$ = 14). **S:** Same-sex pair ($n$ = 10). **D:** Different-sex pair ($n$ = 15). **Male-Male:** Male caregiver with a male youth ($n$ = 3). **Male-Female:** Male caregiver with a female youth ($n$ = 4). **Female-Male:** Female caregiver with a male youth ($n$ = 11). **Female-Female:** Female caregiver with a female youth ($n$ = 7). $^{+}$ $p$ < .01, * $p$ < .05, ** $p$ < .01; *** $p$ < .001.

### C. Supplementary Analyses

#### C-1. Parent–Child Dyadic Neural Similarity and Youth's Emotional Adaptation during Uncertain Aversive Cue

Each component of the emotional adaptation score (depression, anxiety, and ego resilience) was tested individually, and all results remained significant after applying FDR correction. Specifically, higher parent–child similarity in amygdala activation was associated with lower levels of youth depression (B = -0.492, SE = 0.193, 95% CI = [-0.950, -0.165]) and anxiety (B = -0.822, SE = 0.373, 95% CI = [-1.605, -0.116]), as well as higher levels of ego resilience (B = 0.933, SE = 0.393, 95% CI = [0.018, 1.606]). When accounting for age and sex effects, the association between parent–child amygdala activation similarity and youth depression remained significant (B = -0.466, SE = 0.242, 95% CI = [-0.982, -0.034]), while marginal significance was found for youth anxiety (B = -0.751, SE = 0.482, 95% CI = [-1.735, 0.176]) and youth ego resilience (B = 0.888, SE = 0.479, 95% CI = [-0.105, 1.841]).

#### C-2. Parent–Child Dyadic Neural Similarity and Youth's Emotional Adaptation during Regulation of the Uncertain Aversive Cue

Each component of the emotional adaptation score (depression, anxiety, and ego resilience) was tested individually, and significant associations were found for depression and anxiety, but not for ego resilience. Specifically, higher parent–child similarity in amygdala activation was associated with lower levels of youth depression (B = -0.323, SE = 0.172, 95% CI = [-0.730, -0.040]) and lower anxiety (B = -0.447, SE = 0.283, 95% CI = [-1.199, -0.038]), while the association with ego resilience was not significant (B = 0.394, SE = 0.363, 95% CI = [-0.304, 1.154]). When controlled for age and sex effects, the associations between parent-child amygdala activation similarity and anxiety (B = -0.452, SE = 0.295, 95% CI = [-1.225, -0.015]) remained significant, while youth depression (B = -0.306, SE = 0.184, 95% CI = [-0.728, 0.005]) became marginal significant. However, none of the effects survived FDR correction.

#### C-3. Family Relationships and Parent–Child Dyadic Neural Similarity during Uncertain Aversive Cue

Each component of the perceived parent–child relationship score (conflict, cohesion, and identity) was examined individually to assess its specific contribution. Results showed that lower levels of parent–child conflict (B = -0.165, SE = 0.064, 95% CI = [-0.286, -0.039]), and greater sense of family identity (B = 0.144, SE = 0.077, 95% CI = [0.008, 0.314]) was associated with higher parent–child similarity in amygdala activation. However, no significant association was found between parent–child cohesion and parent–child amygdala similarity (B = 0.026, SE = 0.074, 95% CI = [-0.117, 0.173]). After FDR correction, only the effect of parent–child conflict remained significant, while the association with family identity did not survive correction. When controlling for age and sex, effects regarding parent–child conflict remained significant (B = -0.173, SE = 0.084, 95% CI = [-0.328, -0.004]), while effects regarding family identity became marginal significant (B = 0.141, SE = 0.085, 95% CI = [-0.010, 0.323]).

#### C-4. Parent–Child Conflict as a Predictor in the Mediation Model

#### Uncertain Aversive Cue

Specifically, Model 1 (Figure S1A) was conducted to examine whether parent–child neural similarity during uncertainty-induced affective processing mediated the association between youth-perceived parent–child conflict and their emotional adaptation. A significant indirect effect was found (B = -0.493, SE = 0.395, 95% CI = [-1.534, -0.060]); while the direct effect of youth-perceived parent–child conflict on emotional adaptation was not significant (B = -

0.780, SE = 0.834, 95% CI = [-2.404, 0.868]), suggesting that parent–child neural similarity fully mediated this association. The indirect effect accounted for 61.27% of the total effect (B = -1.273, SE = 0.658, 95% CI = [-2.799, 0.229]). Model 2 examined another mediation pathway, with neural similarity as the independent variable and perceived parent–child conflict as the mediator. This model was also not significant in indirect effect (B = 1.092, SE = 1.091, 95% CI = [-0.782, 3.901]), suggesting no evidence of mediation in this direction (Figure S1B).

Figure S1. Mediation Models Examining the Role of Parent–Child Neural Similarity in Amygdala and Youth-Perceived Parent–Child Conflict in Predicting Youths' Emotional Adaptation.

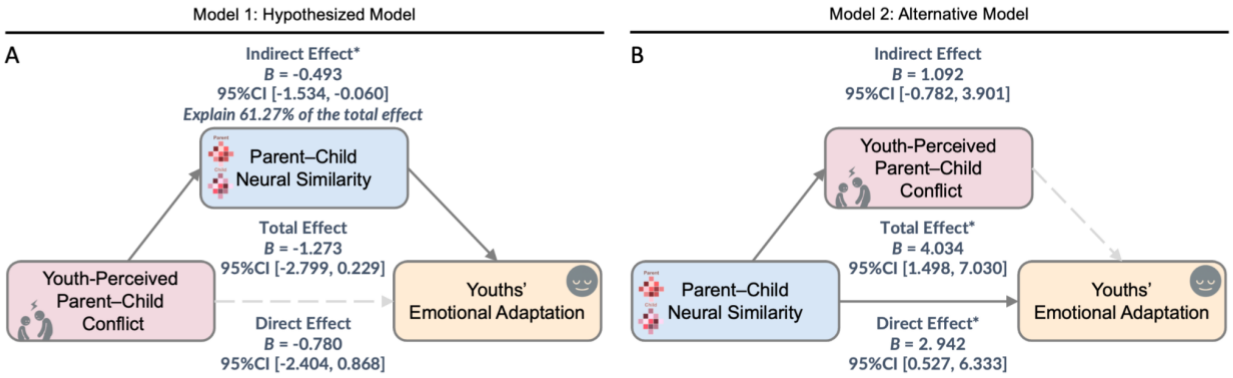


(A) Hypothesized mediation model: Youth-perceived parent–child conflict is negatively associated with parent–child neural similarity in the amygdala during the uncertain aversive cue phase, which in turn is associated with better youth emotional adaptation. The indirect effect was significant, accounting for 61.27% of the total effect. (B) Alternative model: Parent–child neural similarity predicts youth-perceived parent–child conflict, which then predicts emotional adaptation. The indirect path was not significant. * denotes significant effects (95% Confidence intervals does not include zero). Shaded paths indicate non-significant effects.

**D. Amygdala Connectivity Map Associated with Parent–Child Neural Similarity During the Uncertain Cue Phase**

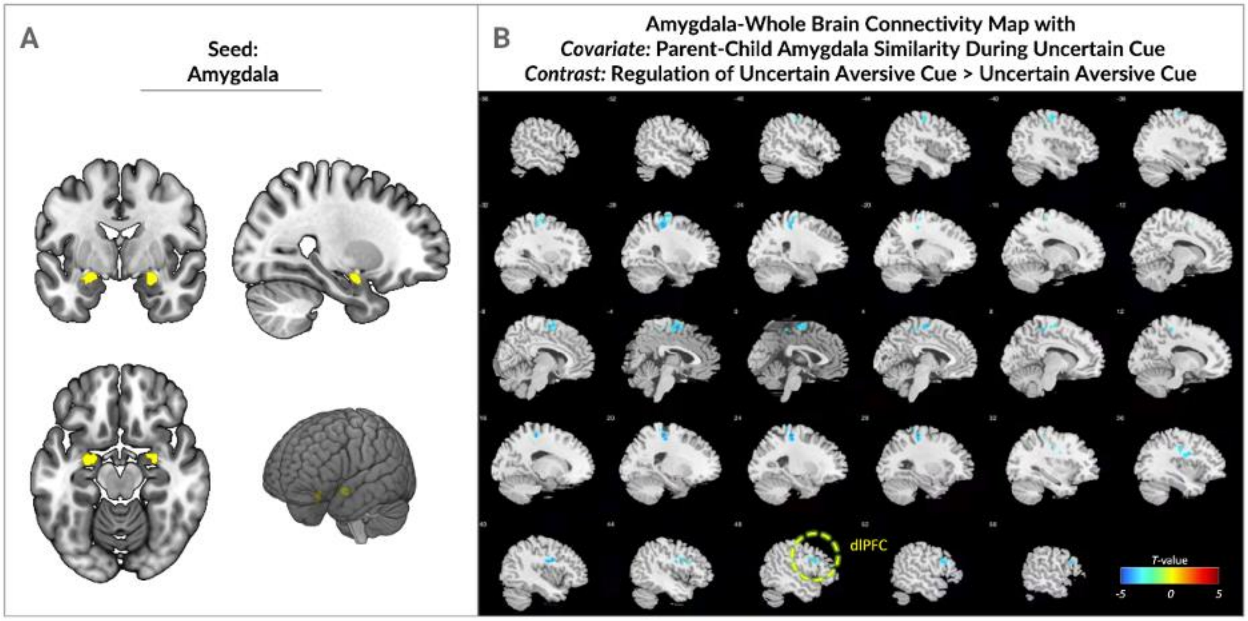


Figure S2: (A) The amygdala seed region used for neural similarity calculation and connectivity analysis is highlighted in yellow. (B) Whole-brain connectivity results from the amygdala seed are shown, with regions exhibiting significant associations with parent–child amygdala similarity during the uncertain cue phase. Increased connectivity between the amygdala and the dorsolateral prefrontal cortex (dlPFC) is circled. The color bar represents *T*-values, ranging from -5 (blue) to +5 (red), indicating the strength and direction of the associations.

**E. Amygdala Connectivity Map Associated with Parent–Child Neural Similarity During the Regulation of Uncertain Cue Phase**

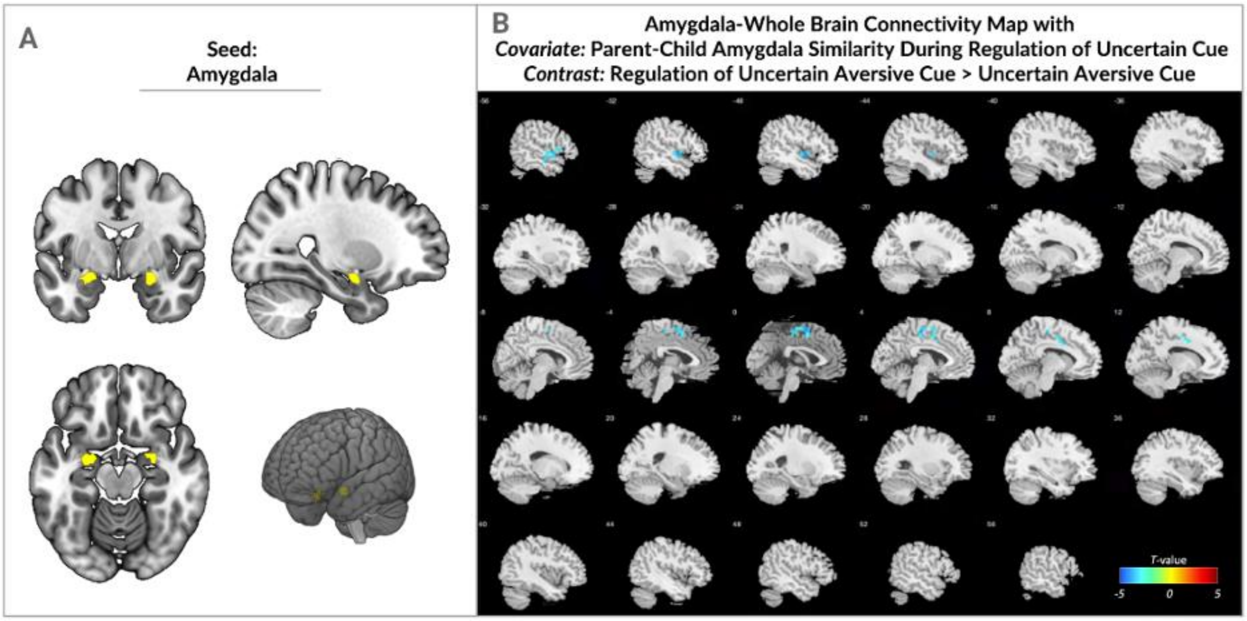


Figure S3: (A) The amygdala seed region used for connectivity analysis is highlighted in yellow. (B) Whole-brain connectivity results from the amygdala seed are shown, highlighting regions that exhibited significant associations with parent–child amygdala similarity during the regulation phase of the uncertain cue. No prefrontal regions were identified as significant in this analysis. The color bar represents *T*-values, ranging from -5 (blue) to +5 (red), indicating the strength and direction of the associations.
